# Population structure of *Borrelia garinii* from *Ixodes uriae* collected in seabird colonies of the northwestern Atlantic Ocean

**DOI:** 10.1101/296319

**Authors:** Hannah J. Munro, Nicholas H. Ogden, Samir Mechai, L. Robbin Lindsay, Gregory J. Robertson, Hugh Whitney, Andrew S. Lang

## Abstract

The occurrence of *Borrelia garinii* in seabird ticks, *Ixodes uriae*, associated with different species of colonial seabirds has been studied since the early 1990s. Research on the population structure of this bacterium in ticks from seabird colonies in the northeastern Atlantic Ocean has revealed admixture between marine and terrestrial tick populations. We studied *B. garinii* population structure in *I. uriae* collected from seabird colonies in the northwestern Atlantic Ocean, in Newfoundland and Labrador, Canada. We applied a multi-locus sequence typing (MLST) scheme to *B. garinii* found in ticks from four species of seabirds. The *B. garinii* strains found in this seabird colony ecosystem were diverse. Some were very similar to strains from Asia and Europe, including some obtained from human clinical samples, while others formed a divergent group specific to this region of the Atlantic Ocean.

**Importance:** This study provides the first *B. garinii* sequences from North American seabird ticks that were characterized using an MLST approach. This revealed new MLST sequence types and alleles, enhancing our knowledge of *B. garinii* diversity. Our findings highlight the genetic complexity of *B. garinii* circulating among seabird ticks and their avian hosts but also demonstrate surprisingly close connections between *B. garinii* in this ecosystem and terrestrial sources in Eurasia. Genetic similarities among *B. garinii* from seabird ticks and humans indicate the possibility that *B. garinii* circulating within seabird tick-avian host transmission cycles could directly, or indirectly via connectivity with terrestrial transmission cycles, have consequences for human health.

## Introduction

*Borrelia burgdorferi* sensu lato (s.l.) is a bacterial species complex that includes the causative agents of Lyme disease, the most common vector-borne disease in the Northern Hemisphere. In North America, *B. burgdorferi* sensu stricto (s.s.) is the only genospecies known to cause Lyme disease in humans to-date, even though several other members of the species complex have been isolated from ticks in the family Ixodidae on the continent (1–4). *Borrelia burgdorferi* is transmitted to humans in North America by *Ixodes scapularis* (in eastern, central, and southern regions) and *I. pacificus* (in western, particularly Pacific coastal, areas). In Eurasia, *B. afzelii*, *B. garinii*, *B. burgdorferi* and other *Borrelia* spp. are known to cause Lyme disease in humans (5–7). The main vectors are *I. ricinus* in western Europe and *I. persulcatus* in eastern Europe and Asia. Reservoir hosts vary among the bacterial genospecies, with *B. afzelii* associated with rodents, *B. garinii* associated with birds, and *B. burgdorferi* s.s. a generalist for which both birds and rodents are reservoirs (8).

The transmission cycles of these bacteria, and the risk of human exposure to infected ticks, generally occur in woodland habitats in which ticks can survive during non-parasitic periods of their lifecycle, and where the mammalian and avian hosts for the ticks and bacteria are found (9). However, *B. garinii* was also found in *I. uriae* feeding on Razorbills (*Alca torda*) in the early 1990s in a seabird colony off the coast of Sweden (10). Other seabirds, such as puffins (11), are now also recognized as competent reservoirs of this bacterium, and humans can be infected via *I. uriae* (11). The distribution and prevalence of *B. garinii* in *I. uriae* has now been studied in seabird colonies worldwide, and it has been found associated with a variety of seabird species in both the Northern and Southern Hemispheres (12–15).

The circulation of *B. garinii* in the seabird reservoir is complex (15), spanning a huge geographic range with many possible vertebrate hosts but *I. uriae* as the only known vector species. More genetic diversity has been found in the marine *I. uriae*-*B. garinii* system compared to the terrestrial realm involving *I. ricinus* (16). There is evidence of transhemispheric-scale *B. garinii* movements based on the presence of identical marker gene sequences in both the Northern and Southern Hemispheres (12), shared genetic structure between the Atlantic and Pacific Ocean basins (15), and little apparent genetic population strucutre within these ocean basins (15). Recombination analysis has also demonstrated admixture between the terrestrial and marine genetic pools (15) and it is therefore important to study both the marine and terrestrial *B. garinii* cycles to understand circulation of this bacterium.

The genome of *B. burgdorferi* s.l. consists of a linear chromosome, which carries the genes for cell maintenance and replication, and a large number of linear and circular plasmids, which encode most of the outer surface proteins (Osp) that mediate interactions with hosts and vectors (17). Previously, DNA-DNA hybridization and 23S-5S intergenic spacer (IGS) sequences were used to delineate *Borrelia* species (5, 18). Attempts to classify strains have also utilized 16S-23S intergenic spacer (IGS) sequences (19, 20) and the plasmid-encoded *ospA* and *ospC* genes (21, 22). Multi-locus sequence typing (MLST), using core housekeeping genes, has become more widely used (19, 23–25) as this allows for analysis at multiple genetic levels, from delineation of species (26) to exploration of population structure (24).

Previous studies have documented *B. garinii* in seabird colonies and shown genetic evidence for linkage between strains in terrestrial and marine environments (15, 27), but samples from North American seabird colonies have never been included. Here we characterize the population structure of *B. garinii* circulating within seabird colonies in the northwestern Atlantic Ocean. To our knowledge, this represents the first sequence-based study from this region. Using an MLST scheme (23) that is currently considered the gold-standard and which has been applied to multiple *Borrelia* species and strains worldwide we examined the population structure of *B. garinii* in *I. uriae* in this marine ecosystem and how it relates to *B. garinii* found throughout Eurasia.

## Results

### Identification of *B. garinii* sequence types

A total of 20 *B. garinii*-positive *I. uriae* collected from four seabird colonies in the northwestern Atlantic Ocean (Figure 1) were used in this study. Nucleotide sequences for eight MLST loci were determined and used to define sequence types (STs). This produced 12 different STs, 10 of which were novel (assigned ST numbers 684 and 686-694). The novel STs contained 26 new alleles and 18 that already existed within the pubMLST database (Supplementary Table S1). Only two previously identified STs were found: ST244 and ST575. The richness of neither STs nor alleles reached saturation in a species accumulation analysis (Supplementary Figure S1), indicating that increased sampling would result in more unique STs and alleles within this population.

**Figure 1.**
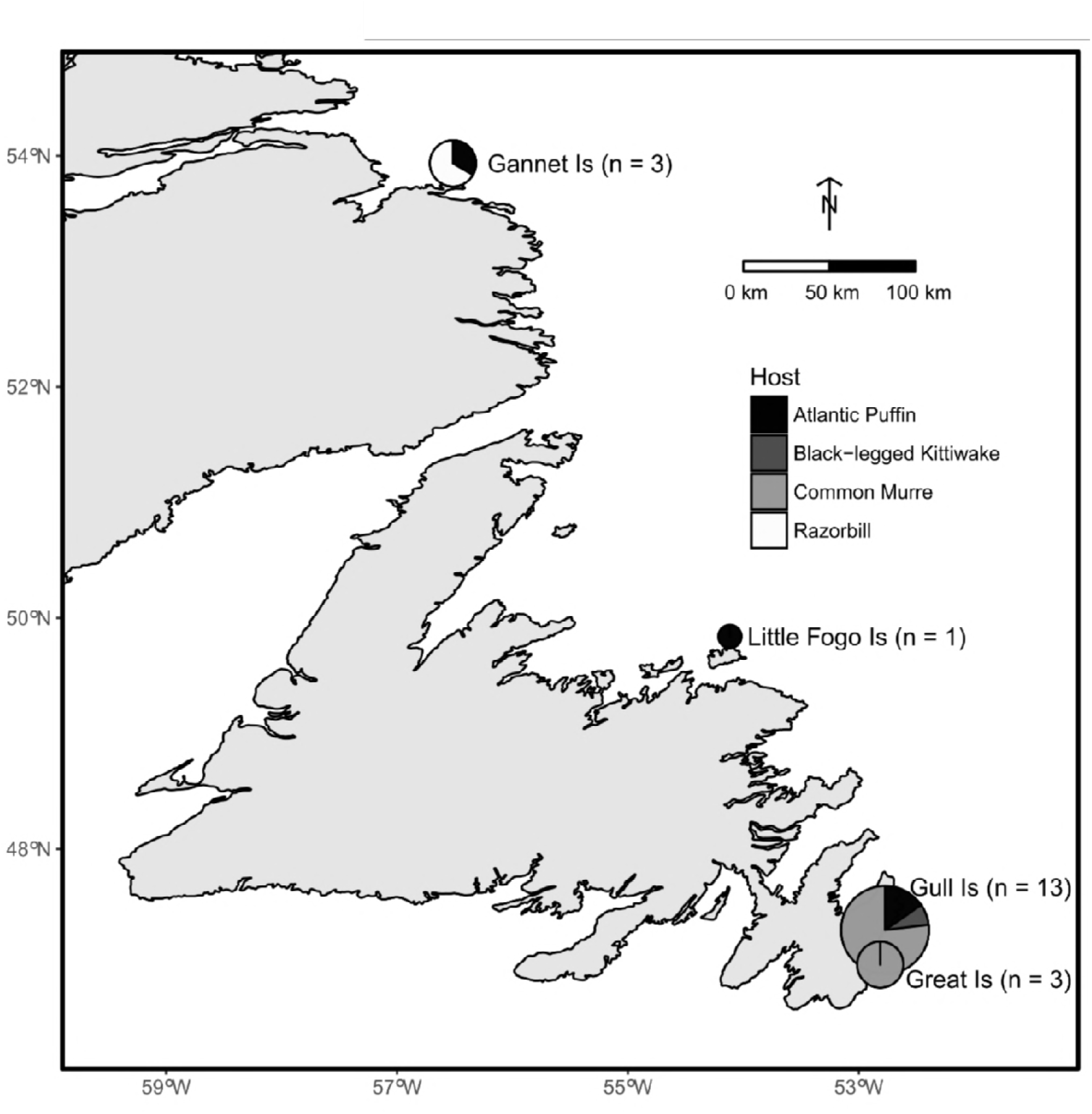
Geographic locations and avian host species compositions for *Ixodes uriae* sources of *Borrelia garinii* sequences used in this study. The proportions of the different host species are denoted in the pie charts and the numbers of *I. uriae* from each location are in brackets. The map was made using the package “maps” in R (67).

The 12 identified STs were distributed across the four colonies and were identified in ticks collected from four seabird hosts (Table 1). The two previously described STs were found on Gull Island (ST244 and ST575) and Little Fogo Islands (ST575), in ticks collected near (and presumed to have fed on) breeding Common Murres (*Uria aalge*) and Atlantic Puffins (*Fratercula arctica*), respectively. Novel STs were found at all colonies and associated with all seabird species investigated. The richness (the number of STs per location or seabird host) did not differ between Gull Island and other locations or between Common Murre and other hosts. Of the 12 STs identified, unique STs (those not found at another location or associated with another host) were found at each location except for Little Fogo Islands, and were associated with both Common Murres and other seabird hosts (Table 1, Supplementary Table S2, Supplementary Figure S2). The proportion of samples carrying unique STs did not differ between Gull Island and other locations (Fisher exact test, χ^2^ = 0.016, p = 0.90), or between Common Murre and other seabird hosts (Fisher exact test, χ^2^ = 0.730, p = 0.39). The relationship between ST richness based on geographic location and host were not independent, with the majority of the samples from Gull Island originating from Common Murres (10 out of 13) whereas at other locations the distribution of ticks among presumed host species was more even (3 out of 7).

**Table 1.** Sequence types (STs) by colony and tick host.

|  | Number<br>of<br>samples | Number<br>of STs | Number of<br>new STs | Number (proportion)<br>of samples carrying<br>unique STs |
| --- | --- | --- | --- | --- |
| Colony |  |  |  |  |
| Gull Island | 13 | 9 | 2 | 6 (0.46) |
| Other | 7 | 6 | 1 | 3 (0.43) |
| Great Island | 3 | 2 | 0 | 1 (0.33) |
| Little Fogo Islands | 1 | 1 | 1 | 0 |
| Gannet Islands | 3 | 3 | 0 | 2 (0.67) |
| Host |  |  |  |  |
| Common Murre | 12 | 9 | 2 | 7 (0.58) |
| Other | 8 | 5 | 1 | 3 (0.38) |
| Atlantic Puffin | 5 | 4 | 1 | 1 (0.20) |
| Black-legged<br>Kittiwake | 1 | 1 | 0 | 0 |
| Razorbill | 2 | 2 | 0 | 1 (0.50) |

### Phylogenetic relationships

The sequences found in our study were phylogenetically diverse, with two sequences branching alone and the others falling into three multi-sequence clades (Figure 2, Supplementary Table S2). Two of these clades, C1 and C4, contained sequences from multiple locations and different host bird species. Each of these clades contained one of the two previously identified STs and clade C4 also contained additional reference sequences from Europe. The third multi-sample clade, C5, contained sequences exclusively from Common Murres on Gull Island and no reference sequences. One of the lone sequences, C2, was basal to clade C1, sharing 99.8% nucleic acid identity with sequences in C1 but differing at every locus with the closest pre-existing ST. The second lone sequence, C3, was basal to a clade of sequences from Europe, with which it shared no alleles at 100% identity.

**Figure 2.**
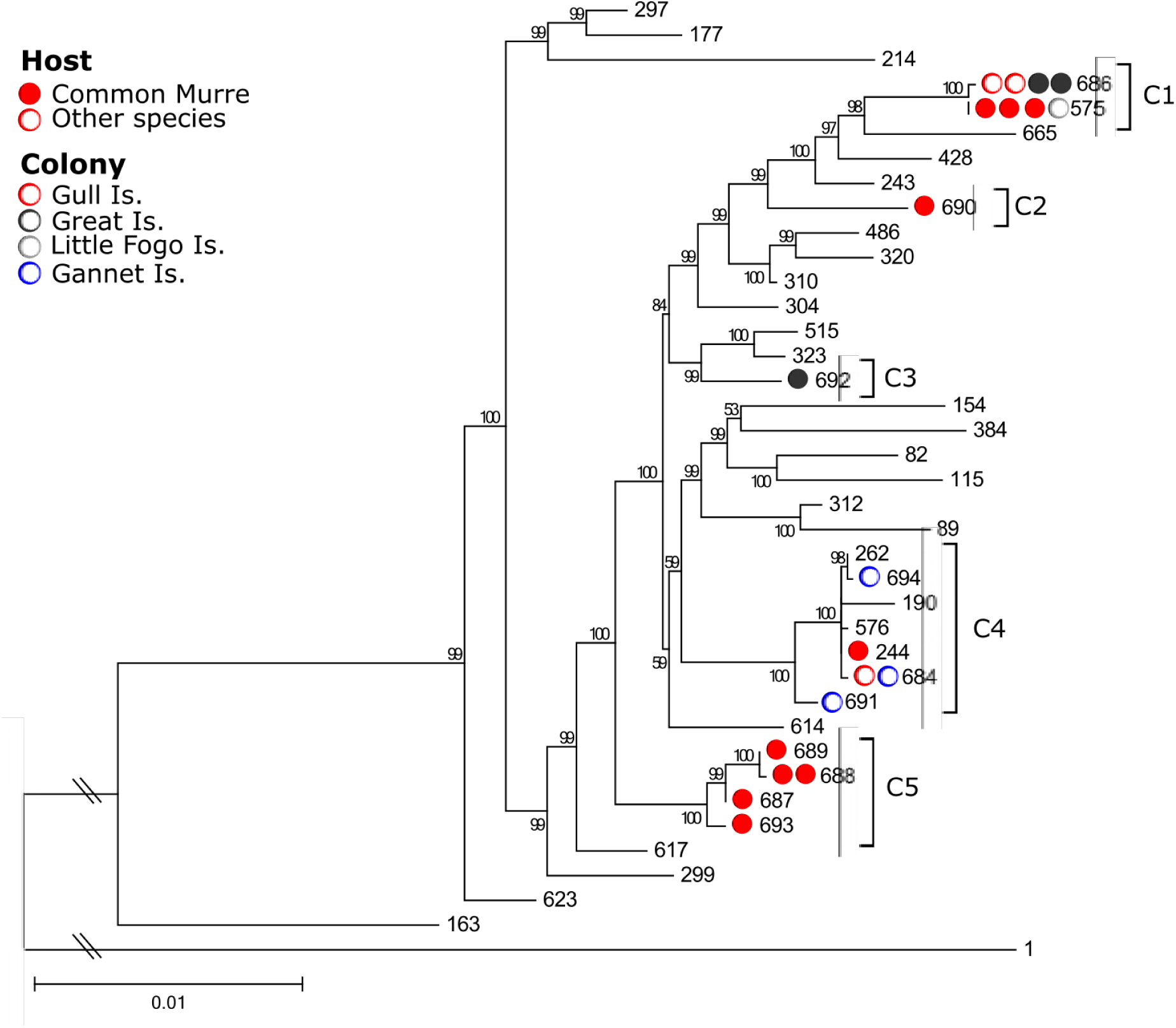
Phylogenetic analysis of *B. garinii* sequences in *I. uriae* from seabirds in the northwestern Atlantic Ocean. The maximum likelihood phylogeny was constructed using PhyML for eight concatenated MLST genes (*clpA, clpX, nifS, pepX, pyrG, recG, rplB, and uvrA*). Labels are sequence types (STs) from the pubMLST database. Sequences from this study are denoted with circles, where colors indicate colony, filled circles represent samples from Common Murres, and empty circles are all other bird species. *Borrelia burgdorferi* s.s was used as the outgroup, labeled as “1”. Numbers at branch nodes represent support based on aBayes and the scale bar represents the number of substitutions per site. The five branches/clades with sequences from this study are denoted C1 through C5.

### Population genetic structure

Pairwise F_ST_ values (Table 2) indicated genetic differentiation and population structuring among localities and tick host species. Comparison of STs from Little Fogo Islands and Great Island showed the highest genetic differentiation values (F_ST_ = 0.733, p < 0.01). Lesser, but still significant, genetic differentiation was found between STs from Little Fogo Islands and Gull Island (F_ST_ = 0.228, p < 0.01). The Gannet Islands showed significant differentiation from Gull Island (F_ST_ = 0.049, p < 0.01) but less than found between Little Fogo Islands and the islands in Witless Bay (Gull and Great Islands). Atlantic Puffin STs showed differentiation from all other species, with the largest value for genetic differentiation being from Razorbills (F_ST_ = 0.695, p < 0.01) and the least with Common Murres (F_ST_ = 0.248, p < 0.01). Genetic differentiation varied more among host species than geographic localities/colonies.

**Table 2.** Matrix of pairwise F_ST_ values of STs between colonies and bird species, with 99% CI.

| Location | Gannet Islands | Great Island | Gull Island |
| --- | --- | --- | --- |
| Great Island | 0 (0-0) |  |  |
| Gull Island | 0.05 (0.01-0.08)* | 0 (0-0) |  |
| Little Fogo Islands | 0.18 (0-0.32) | 0.73 (0.32-0.17)* | 0.23 (0.07-0.33)* |
| Host | Atlantic Puffin | Black-legged Kittiwake | Common Murre |
| Black-legged Kittiwake | 0.49 (0.28-0.65)* |  |  |
| Common Murre | 0.25 (0.11-0.35)* | 0.01 (0-0.05) |  |
| Razorbill | 0.70 (0.28-1)* | 0 (0-0) | 0 (0-0) |

\*Significant comparisons ( $p < 0.01$ )

We performed an eBURST analysis with all 130 *B. garinii* STs, which revealed that the samples clustered into 21 clonal complexes (using the single-locus variant criterion; SLV) and 63 singletons with eight possible founders. The 12 STs found in this study clustered into four clonal complexes when either SLVs or both SLVs and double-locus variants (DLVs) were included, including three singletons (Figure 3, Supplementary Table S2). In this analysis only one clonal complex had an inferred founder, ST244, previously identified in tick and human samples from Germany, Russia, and the UK.

**Figure 3.**
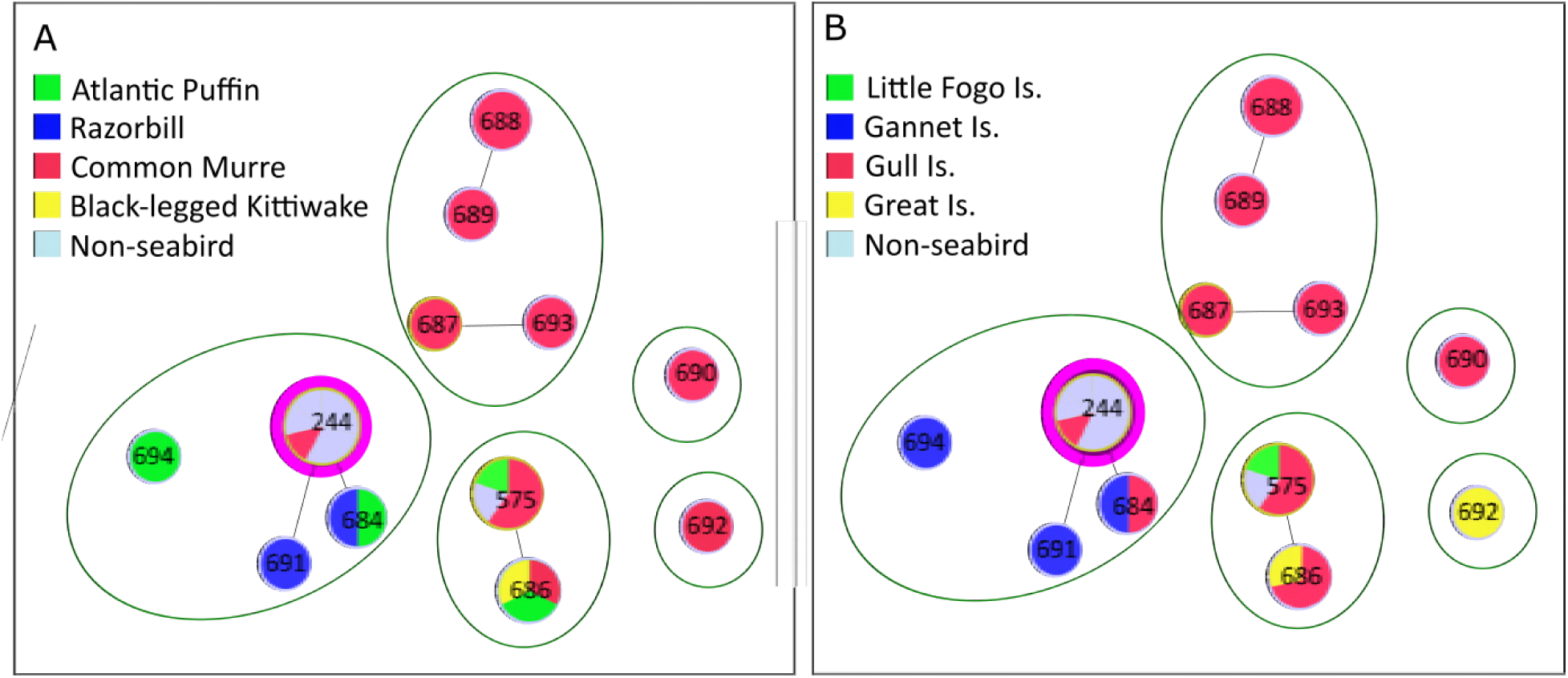
goeBURST network of the 12 sequences types (STs) of *B. garinii* from this study. The STs are highlighted by seabird host (A) and colony (B). The lines denote connections within clonal complexes. The sizes of the circles are proportional to number of samples in the STs. The dark green circles denote BAPS clusters. Inferred founder STs with > 60% bootstrap support are highlighted in pink. Reference sequences originating from sources other than seabirds are indicated in light blue.

We also performed a Bayesian Analysis of Population Structure (BAPS), which suggested the existence of five subpopulations (Figure 3, Supplementary Table S2) with the highest log marginal likelihood values. These subpopulations showed some geographic structuring, with all STs from the Gannet Islands clustering together with two STs found on Gull Island. There were two subpopulations exclusively from Gull Island, and a single ST found on Great Island formed a solo subpopulation. The final subpopulation contained STs found on Gull Island, Great Island, and Little Fogo Islands. The subpopulations also showed some host structuring, with three subpopulations representing STs only found associated with Common Murres. The remaining two subpopulations contained STs found associated with multiple hosts.

The BAPS using all *B. garinii* STs supported the existence of three subpopulations, with one containing our samples from the northwestern Atlantic region. This subpopulation consisted of samples from across Eurasia and showed no geographic structure. This subpopulation also contained sequences that originated from a range of sources, including various species of ticks and ticks collected from humans.

## Discussion

In this study, *B. garinii* within *I. uriae* collected from seabird colonies of the northwestern Atlantic Ocean were analysed by MLST. This comprehensive genetic analysis of *B. garinii* from North America and this ecological system increases the known genetic diversity of *B. garinii* and contributes to our understanding of this species globally. We determined that there is population structure in *B. garinii*, at both regional and global scales. At the regional scale, sequences show evidence of genetic clustering by both geographic sites and/or seabird hosts. Sequences found in the northwestern Atlantic region do not all cluster together, which might reflect several indepedendent introductions of the bacterium into this region and/or prolonged circulation with diversification over time. There is also similarity of the northwestern Atlantic sequences to those found in terrestrial ticks and clincial samples from humans in Europe, suggesting connectivity with non-marine *B. garinii* transmission cycles (15).

Although other species of *Borrelia* have been found circulating in the *I. uriae*-seabird system, including *B. burgdorferi* s.s., *B. bavariensis*, and *B. lusitaniae*, *B. garinii* is predominant (11, 12, 27, 28). Two STs we found are identical to STs previously identified in Europe. Indeed, one of these STs has a wide geographic range and is represented by six samples in the pubMLST database from the UK, Germany, and Russia, and is associated with diverse sources (e.g., human cerebrospinal fluid, and *I. persulcatus* and *I. ricinus* ticks). We found this ST in a tick collected from a Common Murre on Gull Island. The other previously described ST has only been found in Germany, where it was obtained from a human skin sample collected in 1994.

The high level of sequence identity for samples from North American and Eurasian sources indicates connectivity of *B. garinii* populations across the North Atlantic Ocean. Furthermore, the documentation of mutliple STs in both Eurasia and North America indicate there are frequent movements of the bacterium between these regions. Possible scenarios for movement of the bacterium include transport in infected ticks or in infected birds. Although not impossible, the movement of ticks on seabirds across the Atlantic is unlikely as the period of tick attachment is 4-8 days (29, 30) while it would take many days to cross the Atlantic Ocean and land visits by seabrid species outside the nesting season along the way are unlikely (31, 32). The seabirds studied here generally leave their colonies at the end of the breeding season and spend most of the rest of the year out at sea feeding, with no visits to land before the subsequent breeding season. Therefore, it is more likely that bacteria are moved between colonies in infected birds, epecially if the birds remain persistently infected, as is often the case for mammalian hosts (33), and perhaps for *B. burgdorferi* s.l. in some woodland bird species (34). Adult seabirds have high nest-site fidelity but young adults are known to prospect for new breeding locations, resulting in dispersal of birds, and perhaps *B. garinii*, among colonies (31, 35).

High genetic diversity has been documented in past studies of *B. garinii* in *I. uriae* and seabirds (14–16) and this was also observed in our data. Twelve STs are present in the 20 ticks analyzed, along with many unique alleles. A similarly high level of richness is also seen in Europe (36). In contrast, a much lower richness is observed in *B. burgdorferi* s.s. in North America, with 111 STs identified in 564 samples, although diversity of *B. burgdorferi* s.s does differ among geographic regions (37). *Borrelia garinii* is known to be one of the most heterogeneous of the *Borrelia* species, having both high genetic and antigenic diversity (38). It is likely that the diversity found in our study is only a small snapshot of what actually exists in these seabird-tick ecosystems and more novel STs are likely to be found with further sampling, as was supported by the species accumulation analysis.

The *B. garinii* found in the northwestern Atlantic region show surprising phylogeographic relationships to sequences collected throughout Eurasia, from *I. uriae* in seabird colonies in the eastern Atlantic Ocean, non-marine ecosystems and humans. Our sequences are dispersed throughout the *B. garinii* MLST tree, and some show close relationships with those from throughout Eurasia. This suggests multiple movements of strains and mixing between regions (12, 15, 16, 39). When our samples are examined within the overall *B. garinii* clonal complex structure, they do cluster into the same complex and subpopulation. This reflects the highly clonal nature of this bacterium, with populations existing as clusters of closely related genotypes (or complexes) that are globally distributed and stable over time (40).

At a local level, our data show that a high level of *B. garinii* diversity exists in the northwestern Atlantic seabird colonies, with several independent and divergent clonal groups, consistent with what is found in the eastern Atlantic (15). The distribution of genotypes shows some heterogeneity. One cluster of STs (ST693, ST687, ST688, and ST689) originated solely from Common Murres on Gull Island in both 2012 and 2013. The other two clusters both comprise multiple ticks and originate from two or more colonies and two or more seabird hosts. Additionally, one cluster consists of STs primarily originating from non-Common Murre hosts, with four such ticks giving rise to three STs (ST694, ST684, and ST691) and a single Common Murre tick containing the other ST (ST244) in this cluster.

There was no relationship between ST diversity and geographic location or tick host based on the phylogenetic and BAPS analyses, which may suggest that there are no processes limiting transmission of the bacterium at either geographic or host levels. However, examination of the genetic distances between populations by F_ST_ analysis did show some level of differentiation. Significant genetic differentation was observed across large geographic distances, with the Gannet and Little Fogo Islands different from Great and/or Gull Island in Witless Bay. These sites are approximately 500 and 300 km northwest from Witless Bay, repectively. This pattern may be driven by *I. uriae* population structure, which has been observed among colonies in Iceland and Norway (41–43). Vector-borne pathogens co-occur with their hosts and vectors, and the population genetic structure of hosts and vectors is expected to have a strong driving force on the microbe’s structure (44, 45). Lack of genetic distance between Gull and Great Islands is not suprising as they are within 7 km of each other, share similar seabird species compositions, and would have the easiest opportunities for exchanges of birds, ticks and bacteria.

At the tick host level, genetic differentiation exists between STs found associated with Atlantic Puffins and Black-legged Kittiwakes (*Rissa tridactyla*), Common Murres, and Razorbills. Atlantic Puffins use a distinct breeding habitat, nesting in earthen burrows along grassy slopes (46, 47), whereas the other three species are found along rocky cliffs edges (48–50), or talus slopes (51). Therefore, the differences among bird species might be attributable to population structure at the level of *I. uriae* around their seabird hosts on a local geographic level (52, 53) and this could further drive the large geographic patterns seen. Population subdividisions, like those seen among these seabird species, may act as barriers to gene flow for these bacteria and other pathogens (i.e., multiple niche polymorphism (54)).

Overall, this study has contributed to a broader global understanding of *B. garinii* circulation. There is some evidence of host species associations and differentiation across larger geographic distances, but also connectivity among *B. garinii* found in seabird colonies of the northwestern and northeastern Atlantic Ocean and in humans and non-marine ticks of Eurasia. These connections suggest a complicated circulation system with movement across large geographic scales that we propose is linked to bird migration. More research is needed to determine the mechanism(s) connecting the marine and terrestrial ecosystems.

## Methods

### Ethics

Birds were captured and banded under Environment Canada banding permit 10559. This work was carried out under the guidelines specified by the Canadian Council on Animal Care with approved protocols 11-01-AL, 12-01-AL, 13-01-AL, and 14-01-AL from the Memorial University Institutional Animal Care Committee. Lab work was approved under Biosafety Certificate S-103 from the Memorial University Biosafety Committee. Access to the Witless Bay, Gannet Islands, and Cape St. Mary’s Ecological Reserves was through permits from the Parks and Natural Areas Division of the Newfoundland and Labrador Department of Environment and Conservation.

### Ixodes uriae collection and Borrelia screening

Between 2011 and 2014, *I. uriae* ticks were collected from four seabird colonies in the northwestern Atlantic Ocean region in Newfoundland and Labrador, Canada (Figure 1). Birds were captured for a range of research projects and long-term bird-banding programs. The bodies of birds were examined for ticks with special emphasis on the feet and head as these are areas where *I. uriae* are commonly attached (55–57). All tick life-stages were collected: larva, nymph, and adult (Supplementary Table S1). Ticks were collected directly off birds or from nesting habitat. Ticks on hosts were removed with fine forceps and all ticks were placed in pre-labelled vials in the field and stored at -20°C or -80°C until processed further.

DNA was extracted from ticks using the DNeasy Kit (Qiagen). Samples were identified as *Borrelia*-positive using quantitative polymerase chain reaction (qPCR) targeting a conserved portion of the 23S rDNA (58). Positive samples were subsequently used for PCR amplification of genes used previously for *B. garinii* MLST (23): *clpA, clpX, nifS, pepX, pyrG, recG, rplB*, and *uvrA* (Supplementary Table S3). PCR amplifications were performed according to published protocols (23) using GoTaq (Promega). All PCR products were sequenced using Sanger sequencing technology at The Center for Applied Genomics (Toronto, Ontario). Sequences were visually examined for ambiguities, primer sequences were removed, forward and reverse sequences were aligned, and consensus sequences trimmed to the lengths of reference sequences using Geneious 8 (59). The possibility of mixed infections, indicated by mixed peaks on sequence chromatograms, was noted and data from such samples were not included in subsequent analyses.

All sequences were deposited in the NCBI GenBank database with the accession numbers MF536145-MF536294 and added to the pubMLST database (http://pubmlst.org/borrelia/).

### MLST analysis

Sequences from this study were compared using the pubMLST database functions for sequence query (http://pubmlst.org/borrelia/) with each allele being ascribed a number corresponding to an existing identical allele, or a new number in the case that the allele sequence was new to the database. Submissions for new allele ID numbers or sequence types (STs) were made to the pubMLST database as appropriate. Based on allelic profiles of 8 housekeeping genes, each sample was assigned an existing or new (for sequences with new combinations of alleles or novel alleles) ST number (23, 40). Species accumulation curves were plotted, in R using the package ‘vegan’ (60), to examine the increase in ST/allele richness as more samples are considered. The richness of STs was examined relative to geographic locations and host species. Due to uneven sample distribution across geographic locations and host species, and small sample size for some geographic locations and host species, richness was compared between Common Murres and other seabird species at Gull Island and other locations.

### Phylogenetic analysis

To investigate the phylogenetic relationships among *B. garinii* STs, we used the 12 from this study along with all 130 others found within the pubMLST database, all of which originated from Eurasia. Sequences were aligned using MUSCLE (61). Model selection was performed using JModelTest (62, 63) for each locus and a maximum likelihood tree was produced using PhyML for the concatenated loci (64). Branch support was calculated using a Bayesian-like transformation of the approximate likelihood ratio test (aBayes) because of its high statistical power and calculation speed (65). The number of sequences visualized in the tree was limited to those closely related to ours for easier viewing, and the tree was rooted with *B. burgdorferi* s.s. due to its basal nature relative to *B. garinii* (66).

### Population structure analysis

Using sequence data from the 12 STs from this study, two different pairwise F_ST_ analyses were performed in R (67) using the ‘hierfstat’ package (68) to determine the population structure based on colony of sample collection and seabird host. Genetic distance was computed using F_ST_ as previously described (69). To determine the significance of the F_ST_ value, 10,000 bootstraps were performed, and the level of significance was altered from p < 0.05 by Bonferroni correction to a p < 0.01 to account for multiple pairwise comparisons. Genetic distances between populations based on colonies and seabird hosts were determined on this basis.

To identify clonal clustering of our sequences in relationship to all *B. garinii* STs, related clusters of MLST STs were ‘classified’ into clonal complexes using eBURST v3 (http://eburst.mlst.net/) (70) and goeBURST v1.2.1 (71) and then uploaded into the Phyloviz v2 program (72). This analysis was performed with all *B. garinii* STs in the pubMLST database as of April 2017. These programs are designed for use with MLST data and cluster STs using algorithms on a set of hierarchical rules related to the number of single-locus variants (SLVs), double-locus variants (DLVs; eBURST), and triple-locus variants (TLVs; goeBURST). eBURST uses local optimization and is based on a simple model of clonal expansion and divergence, whereas goeBURST allows for global optimization and the identification of the founder ST among the set of STs, and an extended set of tiebreak rules, which leads to improved graphic representation of clonal complexes relating to the ancestral links among ST components. This analysis provides a global perspective of relationships of new STs and previously described STs, showing founders for the populations and closely related samples based on clonal complexes, as opposed to a phylogeny. Nevertheless, clonal complexes from the MLST analysis and clades on the phylogenetic trees are often concordant (25, 73, 74).

The community structure of the different STs found within the northwestern Atlantic was computed with Bayesian Analysis of Population Structure (BAPS) version 6.0 (75), using clustering with a linked locus module and codon model. Mixture analysis was performed with K values from 1 to 12, and optimal partitions were identified based on maximum log marginal likelihood values. The analysis was repeated with all *B. garinii* STs in the pubMLST database to identify STs from across Eurasia that clustered with STs found in the northwestern Atlantic, with K values from 2 to 20. This provided an understanding of community structure of the samples from this study and how they fit together on a regional scale, as well as on a larger global scale, and it allowed for clonal complexes to be classified into clusters.

## Acknowledgements

This work was supported by the Natural Sciences and Engineering Research Council (NSERC) of Canada (341561 to A.S.L.), the Newfoundland and Labrador Forestry and Agrifoods Agency, the Public Health Agency of Canada, and the Memorial University of Newfoundland School of Graduate Studies (baseline funding to H.J.M.). We thank everyone who helped collect ticks and with all aspects of field work. We thank Marta Canuti and Joost Verhoeven for assistance with the phylogenetic analyses and Gabriele Margos for help with MLST assignments.

## Author contributions statement

H.J.M. performed sample collection, laboratory work, statistical analysis, and manuscript writing. N.H.O. contributed to study design, data interpretation, and manuscript writing. S.M. contributed to data interpretation, statistical analysis, and manuscript writing. L.R.L. contributed to study design and manuscript writing. G.J.R. contributed to sample collection and manuscript editing. H.W. contributed to study design and manuscript writing. A.S.L. contributed to data interpretation and manuscript writing.

